# LncRNA DLEU2 regulates Sirtuins and mitochondrial respiratory chain complex IV: a novel pathway in obesity and DOHaD

**DOI:** 10.1101/2021.05.11.442069

**Authors:** Jian Zhang, Matthew Krivacka Kay, Mahua Choudhury

**Affiliations:** Pharmaceutical Sciences, Texas A & M Health Science Center

## Abstract

Long noncoding RNAs (lncRNAs) are commonly dysregulated in cancer but are mostly unknown for roles in metabolic regulation. Sirtuins, an epigenetic modulator class, regulate metabolic pathways. However, how sirtuins are regulated via lncRNA is unknown. In our findings, DLEU2, a lncRNA involved in primarily blood cancers, and sirtuins were both significantly decreased in the livers of high-fat high-fructose diet (HFD-HF) fed male mouse offspring (F1) whose mothers (F0) were either fed chow diet or HFD-HF during reproductive and pregnancy windows. Confirming this connection, upon silencing DLEU2, transcription levels of SIRT1 through 6 and translational levels of SIRT1, 3, 5, and 6 were significantly downregulated. Knockdown of DLEU2 also significantly decreased the protein level of cytochrome-c oxidase (complex IV, MTCO1) without altering other mitochondrial complexes, leading to increased reactive oxygen species production. Interestingly, in F1 livers, the protein level of MTCO1 was also significantly decreased under an HFD-HF diet or even under chow diet if the mother was exposed to HFD-HF. Our findings therefore reveal for the first time that a lncRNA can regulate sirtuins and a specific mitochondrial complex, furthermore suggesting a potential role of DLEU2 in metabolic disorders over one or more generations.

**HIGHLIGHTS:** Maternal diet can modulate hepatic lncRNA DLEU2 and its downstream regulators in offspring

DLEU2 is required for specific sirtuin regulation and mitochondrial respiration chain complex IV expression

Offspring liver depleted of DLEU2 show decreased mitochondrial respiration chain complex IV and specific sirtuins at transcriptional and translational levels

## INTRODUCTION

Obesity has emerged as the leading medical disorder of the 21st century with an ever increasing incidence throughout the world (Ng et al., 2014). Several factors have been pinpointed for this increase in obesity; for example, a sedentary lifestyle and improper dietary habits, including a Western diet (Soro-Arnaiz et al., 2016). Western diets, in particular those high in carbohydrates and fats, have led to a procession of obesity-related disorders, including type 2 diabetes, dysglycemia, and metabolic syndrome (Taylor and Poston, 2007). This obesity epidemic is even more severe than at first appearance, as there is increasing evidence showing that exposure to a Western diet to the mother increases the risks of offspring developing features of metabolic syndrome as well, known as the “Developmental Origins of Health and Disease” (DOHaD) theory (Godfrey, 1998; Jones et al., 2009; King, 2006; Samuelsson et al., 2008; Yajnik and Deshmukh, 2008). As the impacts of obesity can pass from one generation to the next, scientists are now investigating how and why these signals for metabolic dysregulation can be inherited. Epigenetic inheritance can play a broad role in enacting these scenarios (Choudhury and Friedman, 2011; King and Skinner, 2020; Li, 2018).

Epigenetics involves the heritable transmission of phenotype changes which do not involve alterations in the DNA sequence (Dupont et al., 2009). These transmissible changes can be through long non-coding RNAs (lncRNAs), which are non-coding RNA fragments known to be more than 200 nucleotides long (An et al., 2017; Ma et al., 2013; Perkel, 2013). lncRNAs were originally considered irrelevant transcriptional noise, but have since been shown to play key roles in both physiological and pathological regulatory pathways (Mazo et al., 2007; Rinn et al., 2007). DLEU2, short for *Deleted in leukemia 2*, is a recently discovered lncRNA which has been shown to play an important role in multiple cancers, including blood (Klein et al., 2010), renal (Lv et al., 2017), and pancreatic (Xu et al., 2019). DLEU2 lncRNA also serves as the intronic site of tumor suppressor microRNAs (miRNAs) miR-15a and miR-16-1, which play downstream roles in controlling cell cycling functions (Lerner et al., 2009). Of considerable note, miR-15a and miR-16-1 have been shown to be involved in metabolic regulation, including insulin signaling and PI3K-Akt pathways (Kwon et al., 2014). However, no connections have been drawn between the lncRNA DLEU2, metabolic disorders, and obesity as of yet.

Epigenetic regulation can also be applied through histone and protein molecular tagging, such as lysine acetylation/deacetylation of terminal residues, to control gene accessibility and expression (Choudhary et al., 2009; Hirschey et al., 2011; Kendrick et al., 2011; Milazzo et al., 2020). Sirtuins, a seven-membered family (SIRT1-7) of epigenetic regulators, located in different subcellular regions, are grouped as the class III histone deacetylases (HDACs) (Feldman et al., 2012). The sirtuins (SIRTs) are separated by their activities, with SIRT1, 2, 6, and 7 located in the nucleus and SIRT 3, 4, and 5 located in the mitochondria (Feldman et al., 2012). Our group and others show that SIRTs display crucial roles in a variety of key metabolic pathways, including energy metabolism, stress response, fatty acid oxidation, triglyceride pathways, inflammation, and mitochondrial regulation (Borengasser et al., 2011; Choudhury et al., 2011; Kendrick et al., 2011; Suter et al., 2012; Zhang et al., 2015). SIRTs thus participate in a variety of aspects, controlling metabolism and stress throughout the body. There are no reports yet connecting all these sirtuin regulations with any specific lncRNA. Interestingly, high-Myc state repressed DLEU2 and Myc binding site was shown to be associated with DLEU2’s promoter region (Chang et al., 2008). While no direct pathways have been implicated, interplay has been shown between sirtuins and Myc cancer-related metabolic reprogramming (Zwaans and Lombard, 2014). Recent evidence has also shown that some HDACs inhibitors, such as trichostatin A (TSA), upregulate DLEU2 and miR-15a/16-1, in lung cancer cells (Chen et al., 2013). However, though cancer links have been discovered, to our knowledge no studies have been performed linking any lncRNAs with sirtuins, especially via dietary factors. Herein we hypothesize DLEU2 may regulate sirtuins.

Sirtuins have been shown to play a major role in mitochondrial modulation and regulation (Carrico et al., 2018; Hirschey et al., 2011; Nasrin et al., 2010). Mitochondria play key roles in maintaining homeostasis by generating ATP, and performing fatty acid oxidation and oxidative phosphorylation (Lombard et al., 2011). The inner mitochondrial membrane (IMM) contains the electron transport chain complexes I-IV, which generate the electrochemical gradient across the IMM. Defects and alterations in mitochondrial complex expression have been linked to metabolic syndrome and obesity (Ngo et al., 2019; Soro-Arnaiz et al., 2016; Sverdlov et al., 2015). As the mitochondria is directly involved in detoxifying radical oxygen species (ROS) generated from fatty acid oxidation, SIRT3 plays key roles by deacetylating mitochondrial superoxide dismutase (SOD2), inducing SOD2 activity (Qiu et al., 2010), deacetylating FOXO3 for transcription (Tseng et al., 2013), and similarly inducing the glutathione and thioredoxin systems (Kong et al., 2010; Potthast et al., 2017). As such, SIRTs have been identified as vital players in regulating both metabolic and mitochondrial factors in the body, but their links to any lncRNA regulation remain unknown.

In this study, we explore a mechanistic regulation of DLEU2 on sirtuins, the mitochondrial complex, and oxidative stress in liver cells.

## RESULTS

### HFD-HF intake increased body weight and liver weight

To investigate the effect of maternal HFD-HF intake on the body weight (BW) and liver weight (LW) of the offspring, measurements of BW and LW were taken at 20 weeks. Maternal high-fat high fructose intake has no effect on both the BW and LW in F1 pups (Fig. 1 A&B, red vs blue or purple vs green). However, the body and liver weight were significantly increased in pups who were fed HFD-HF compared to CD (Fig. 1A&B, red vs purple or blue vs green, p<0.001, respectively).

**Figure 1.**
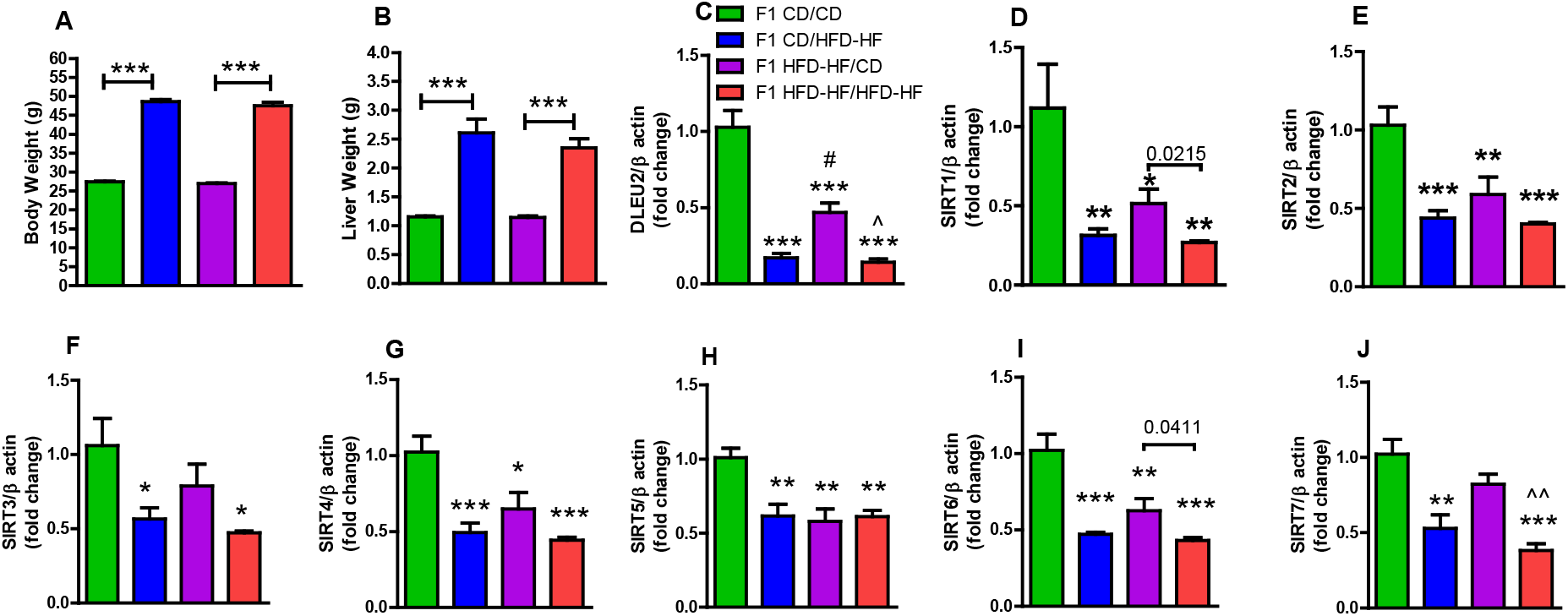
Maternal diet effect on weight of body and liver, and gene expression of DLEU2 and Sirtuins in the livers of F1 male mice on CD and HFD-HF. Body (A) and liver (B) weight of F1 male pups from 4 groups were measured. CD mothers/CD pups, green; CD mothers/HFD-HF pups, blue; HFD-HF mothers/CD pups, purple; HFD-HF mothers/HFD-HF pups, red. Fold change of hepatic RNA expression of C) DLEU2, D) SIRT1, E) SIRT2, F) SIRT3, G) SIRT4, H) SIRT5, I) SIRT6, and J) SIRT7 are shown. Expression was normalized to ß-Actin. n=6 mice/group, one-way ANOVA with Tukey’s post hoc test, *P<0.05, **P<0.01, ***P<0.001 compared to CD/CD. ^#^P<0.05 compared to CD/HFD-HF, ^^P<0.01 compared to HFD-HF/HFD-HF. The p value over comparison line in the figure indicates t-test.

### HFD-HF intake altered hepatic gene expression of DLEU2 and SIRT

Till now, only one article shows the intergenerational effects of HFD on lncRNAs in an array analysis, however, no in depth analysis was carried out on individual level (An et al., 2017). Our study is the first attempt to investigate the intergenerational effect of diet on DLEU2 and sirtuins. In the CD fed F1 offspring, hepatic RNA levels of DLEU2, SIRT1, 2, 4, 5, and 6 were significantly decreased when mother was fed HFD-HF compared to CD fed mother (Fig. 1, C, D, E, G, H, &I, purple vs green P<0.001, P<0.05, P<0.01, P<0.05, P<0.01, and P<0.01, respectively). This showed the intergenerational effects of HFD-HF in our study. However, even in groups with mother on CD (blue and green), gene expression of DLEU2 and all sirtuins were significantly downregulated in offspring fed HFD-HF compared to pups on CD (Fig. 1, blue vs green, One-way ANOVA). These significant effects were not seen for SIRT2, 3, 4, and 5 in offspring with mother on HFD-HF (Fig. 1 E, F, G, &H, red vs purple, One-way ANOVA or t-test). These differences indicate the different effects of maternal diet and pup’s own diet.

### Silencing DLEU2 altered SIRT genes and proteins expression in HepG2 cells

Our *in vivo* study strengthens our hypothesis that DLEU2 may regulate SIRT expression. Therefore, we carried out an investigation of the mechanistic interaction of DLEU2 and Sirtuins in the HepG2 cell line. After DLEU2 was silenced, RNA expression of DLEU2, SIRT1 to 6 were significantly downregulated, but SIRT7 was significantly upregulated (Fig. 2). These results are in line with the above *in vivo* mouse study, except for SIRT7. Consistent with RNA expression, the protein levels of SIRT1, 3, 5, and 6 were decreased after DLEU2 knockdown, however, protein levels of SIRT 2, 4, and 7 were not altered (Fig. 2). These results indicated that DLEU2 may have a direct relationship with SIRT1, 3, 5, and 6.

**Figure 2.**
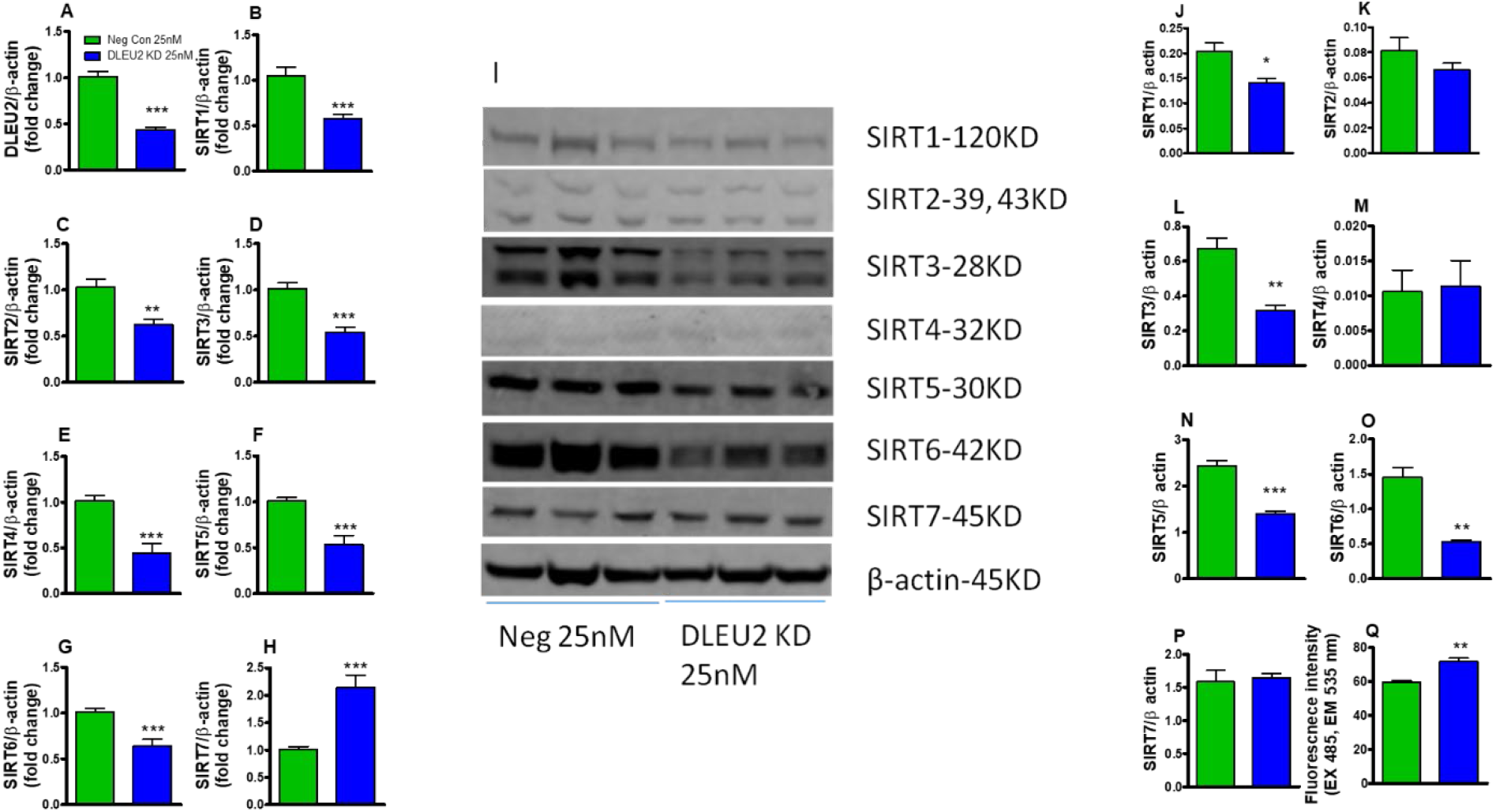
Expressions of DLEU2 and Sirtuins expression and oxidative stress in HepG2 human cancer cell lines after silencing DLEU2. In HepG2 cells, DLEU2 was knocked down for 96 h, and gene expressions of DLEU2 and Sirtuins were analyzed by RT-PCR. RNA levels of A) DLEU2, B) SIRT1, C) SIRT2, D) SIRT3, E) SIRT4, F) SIRT5, G) SIRT6 and H) SIRT7 in DLEU2 knockdown HepG2 cells. Expression was normalized to ß-Actin. n=3. **P<0.01, ***P<0.001 versus negative control. After DLEU2 was knocked down for 96 h, protein expressions of sirtuins were analyzed by western blot. I) ß-Actin was used as a loading control. Densitometry analysis was performed on J) SIRT1, K) SIRT2, L) SIRT3, M) SIRT4, N) SIRT5, O) SIRT6, and P) SIRT7 expression. Expression was normalized to ß-Actin. n=3, *P<0.05, **P<0.01, ***P<0.001 versus negative control. Q) DLEU2 KD increased ROS level in HepG2 cells. Reactive oxygen species (ROS) levels were determined by measuring oxidized dichlorofluorescein (DCF) levels using a 2’, 7’-dichlorofluorescein diacetate (DCFDA) after DLEU2 KD. **P < 0.01 versus negative control; n = 3.

### DLEU2 knockdown decreased Oxphos Complex IV

Sirtuins play roles in mitochondrial function and are involved in oxphos complexes regulation (Verdin et al., 2010), indicating DLEU2 may also play a role in regulating mitochondrial function directly or indirectly. Therefore, mitochondrial complex protein levels were investigated in DLEU2 silenced cells. Interestingly, protein levels of complex IV-MTCO1 was significantly decreased after DLEU2 KD (Fig. 3, A&E, P<0.001) while other mitochondrial complexes were unchanged.

**Figure 3.**
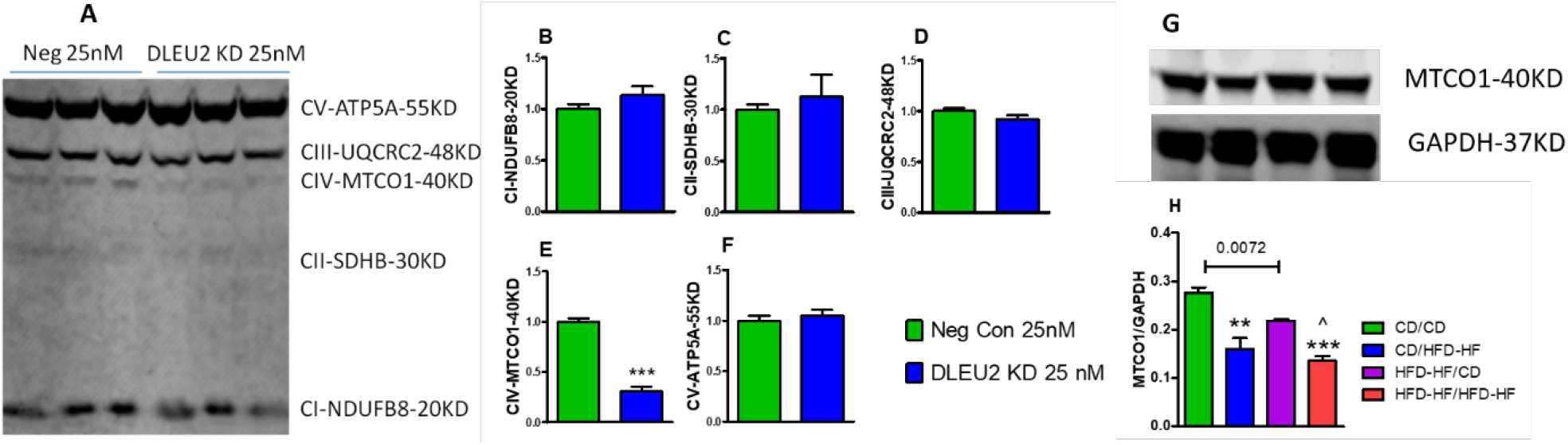
Expressions of OXPHOS in HepG2 human cancer cell line after silencing of DLEU2 and maternal diet effect on protein level of MTCO1 in the livers of F1 male mice on CD and HFD-HF. A) The representative bands by western blot analysis demonstrate the protein levels of OXPHOS complexes B) CI-NDUFB8, C) CII-SDHB, D) CIII-UQCRC2, E) CIV-MTCO1, F) CV-ATP5A after 96 h transfection with Negative control and DLEU2 siRNA. n=6, ***P<0.001 versus negative control. G, H) Protein level of hepatic MTCO1 in F1 liver. Expression was normalized to ß-Actin. n=3 mice/group, one-way ANOVA with Tukey’s post hoc test, **P<0.01, ***P<0.001 compared to CD/CD. ^P<0.05 compared to HFD-HF/HFD-HF. The p value over comparison line in the figure indicates t-test.

### HFD-HF intake in F0 mother altered hepatic protein expression of MTCO1 in F1 generation

Hepatic protein level of MTCO1 in F1 was investigated. Maternal HFD-HF significantly decreased pups’ MTCO1 level compared to CD fed mother (P=0.0072, t-test, purple vs green, Fig. 3 G&H). In F1, HFD-HF feeding also decreased hepatic MTCO1 compared to CD diet (**P<0.01, blue vs green, and ^P<0.05, red vs purple, Fig. 3 G&H).

### Silencing DLEU2 increased oxidative stress

Both sirtuins and mitochondrial complex were shown to contribute in oxidative stress regulation. Oxidative stress may indicate mitochondrial dysfunction (Nita and Grzybowski, 2016). In this study, ROS levels were significantly increased in DLEU2 KD (Fig. 2Q, P < 0.01).

## DISCUSSION

The obesity epidemic is propagated on metabolic syndrome brought about by the worldwide spread of the western diet, with its high fat content and industry-processed sugars and carbohydrates, including fructose, which causes insulin resistance, cardiovascular disease (CVD), and metabolic syndrome-related dysregulation (Abdelmalek MF, 2010; Gotto AM Jr, 2006; Lustig RH, 2016). This problem has become especially acute, as countries that have switched to high fructose corn syrup (HFCS) in their food supply have a 20% higher incidence of diabetes compared to countries that do not use HFCS (Goran MI, 2013). This diet-based premise seems to hold true, as increased weights for both the body and liver were present in our study’s mice feeding on a HFD diet containing high fructose (Fig 1, blue and red bars), as would be expected from multiple related animal studies on HFD with fructose (Charlton et al., 2011; Chukijrungroat et al., 2017; Della Vedova et al., 2016; Liu et al., 2018; Love et al., 2017; Luo et al., 2016; Mastrocola et al., 2018).

It is also well known that alterations in fetal life due to changes in the *in utero* environment may influence the adult onset of metabolic disease as well (Godfrey, 1998; Jones et al., 2009; King, 2006; Samuelsson et al., 2008; Yajnik and Deshmukh, 2008). The DOHaD concept has existed for many years, and has definitively shown that changes to diet and environmental stress during gestation leads to later downstream effects in offspring later in life (Roseboom et al., 2001; Wadhwa et al., 2009). However, our group did not see an *in utero* maternal HFD-HF exposure effect on offspring weight (Fig 1, green and purple vs. blue and red bars). One explanation could be that *in utero* exposure time was not enough to fully affect the offspring’s phenotypes yet at the time of mice sacrifice. However, evidence says that epigenetic changes happen before any symptoms arise. Therefore, it is not unlikely that there may be molecular abnormalities already initiated. As these DOHaD effects can be transmitted epigenetically (Wadhwa et al., 2009), we investigated the various epigenetic changes as these generally happen before actual genetic or phenotype changes which can functionally be reversed (Lacal and Ventura, 2018).

Epigenetic changes are suggested to be introduced from the parents to the offspring via a variety of mechanisms, including DNA methylation, posttranslational histone modification, chromatin remodeling, and non-coding RNA (Horsthemke, 2018; van Otterdijk and Michels, 2016). However, virtually no data are currently available on the role of any specific long non-coding RNA involved in metabolic dysfunction in generational inheritance.

lncRNAs are an under-investigated area in the field of metabolic epigenetics, as most of the current research is being performed in cancer related topics, with a modicum of obesity and diabetes-related subjects. Gonzalez-Rodriguez et al. discovered that heritable growth restriction was associated with changes in lncRNA H19 expression, but such changes to H19 expression were reversible with diet supplementation combating metabolic syndrome (Gonzalez-Rodriguez et al., 2016). As such, we investigated if the lncRNA DLEU2 would have expression changes when exposed to an obesity-causing diet regimen from mother to offspring. The change of DLEU2 in offspring under different diet was also investigated. More importantly, there are virtually no studies connecting DLEU2 with metabolic diseases. DLEU2 expression was found to be significantly decreased in the livers of offspring mice in a likewise manner, with more than 70% downregulated DLEU2 expression in offspring on a HFD-HF compared to on a CD (Fig 1C, >80% down blue vs green and, >70% down red vs purple bars). These specific lncRNA responses to diets led us to investigate other epigenetic modulators that might be regulated by DLEU2. A study by Chen et al. showed that the expression of DLEU2 was found to be increased by the histone deacetylase (HDAC) inhibitor trichostatin A (TSA) in non-small cell lung cancer (Chen et al., 2013). This related finding triggered our interest in searching for a specific DLEU2-HDAC relationship, specifically sirtuin interactions, as we and others have previously established this HDACIII group’s role in metabolic diseases (Choudhury et al., 2011; Kendrick et al., 2011; Zhang and Choudhury, 2017; Zhang et al., 2015).

While the mice offspring exposed to western diet showed significant downregulation of sirtuins (Figure 1D-J, blue and red bars), we found that a maternal HFD-HF decreased hepatic sirtuin expression in liver of offspring. (Figure 1D, E, G, H, and I, purple bars). This result is in support of our previous findings, as our non-human primate study showed that a maternal HFD decreased the hepatic expression of Sirt1 in offspring (Suter et al., 2012). We then followed-up with a mechanistic experiment, where DLEU2 was knocked down in HepG2 cells, to reveal a direct connection between lncRNA and Sirts, as the majority of the sirtuins were downregulated at the transcriptional (Fig 2A-G) and at the translational (Fig 3I, J, L, N, and O) levels. Silencing of DLEU2 *in vitro* had similar genes effect as *in vivo* HFD-HF exposure except SIRT7 (Fig 1J and Fig 2H). This effect was diminished though at the transcriptional level for SIRT 7 as well as SIRT2 and 4 (Fig 2 K, M, and P). Of note, SIRT7 has shown the opposite role compared to SIRT1 in adipogenesis (Bober et al., 2012; Fang et al., 2017).

Overall, we can speculate that there might be other regulators which could try to normalize a DLEU2 effect in a whole body system, however, DLEU2’s effects on SIRT1, 3, 5, and 6 showed similar effects at the transcription and translational levels both *in vivo* and *in vitro* (Fig 2L and N). Interestingly, two of these sirtuins (3 and 5) are localized to the mitochondria (Michishita et al., 2005), and all of these sirtuins are linked to oxidative stress (Bause and Haigis, 2013; Cheng et al., 2016; Liu et al., 2013; Salminen et al., 2013; Singh et al., 2018). SIRT3 has been known to be associated with the electron transfer chain components, including controlling acetylation of complex IV (Kendrick et al., 2011), β-oxidation of fatty acids (Hirschey et al., 2010), and extra-mitochondrial PGC-1α co-activation for mitochondrial biogenesis (Jeninga et al., 2010), with SIRT5 involved in the urea cycle for ammonia detoxification and disposal (Nakagawa et al., 2009). Therefore, we hypothesized that DLEU2 may be the non-coding epigenetic regulator which is carrying out these downstream changes in the mitochondrial complex while modulating oxidative stress.

Fulfilling our expectations, mitochondrial defects were discovered in the form of a significant mitochondrial complex IV reduction from the DLEU2 knockdown and F1 liver, leading to increased oxidative stress (Fig. 2Q & 3). We have previously shown that in *SIRT3*^*-/-*^mice, exposure to a HFD reduced activity of mitochondrial complexes III and IV (Kendrick et al., 2011). These results matched other studies showing higher levels of production of mitochondrial ROS under stress-inducing conditions (Kim et al., 2010; Sundaresan et al., 2009). Potthast et al discovered the inhibition of situins 1, 3 and 4 activity, protein and transcriptional levels in fibroblasts from patients with mitochondrial COX IV-deficiency (Potthast et al., 2017). Our result is specifically novel as DLEU2 is regulating only one mitochondrial complex-therefore a specific targeted regulation will be investigated in our future obesity study using knockout and overexpression mice.

In conclusion, a western high fat high fructose diet disturbs the hepatic regulation of an epigenetic factor DLEU2, potentially leading to metabolic disorders in the same generation or in the next generation (Fig. 4). DLEU2 directly regulates most members of sirtuins and could be a new target for modulation for remedying metabolic dysfunction, reducing oxidative stress. More *in vivo* studies with knockout mouse models are needed to discover the function of DLEU2 in metabolic regulation pathways and also in transgenerational inheritance, especially in the liver or other metabolic tissues.

**Figure 4.**
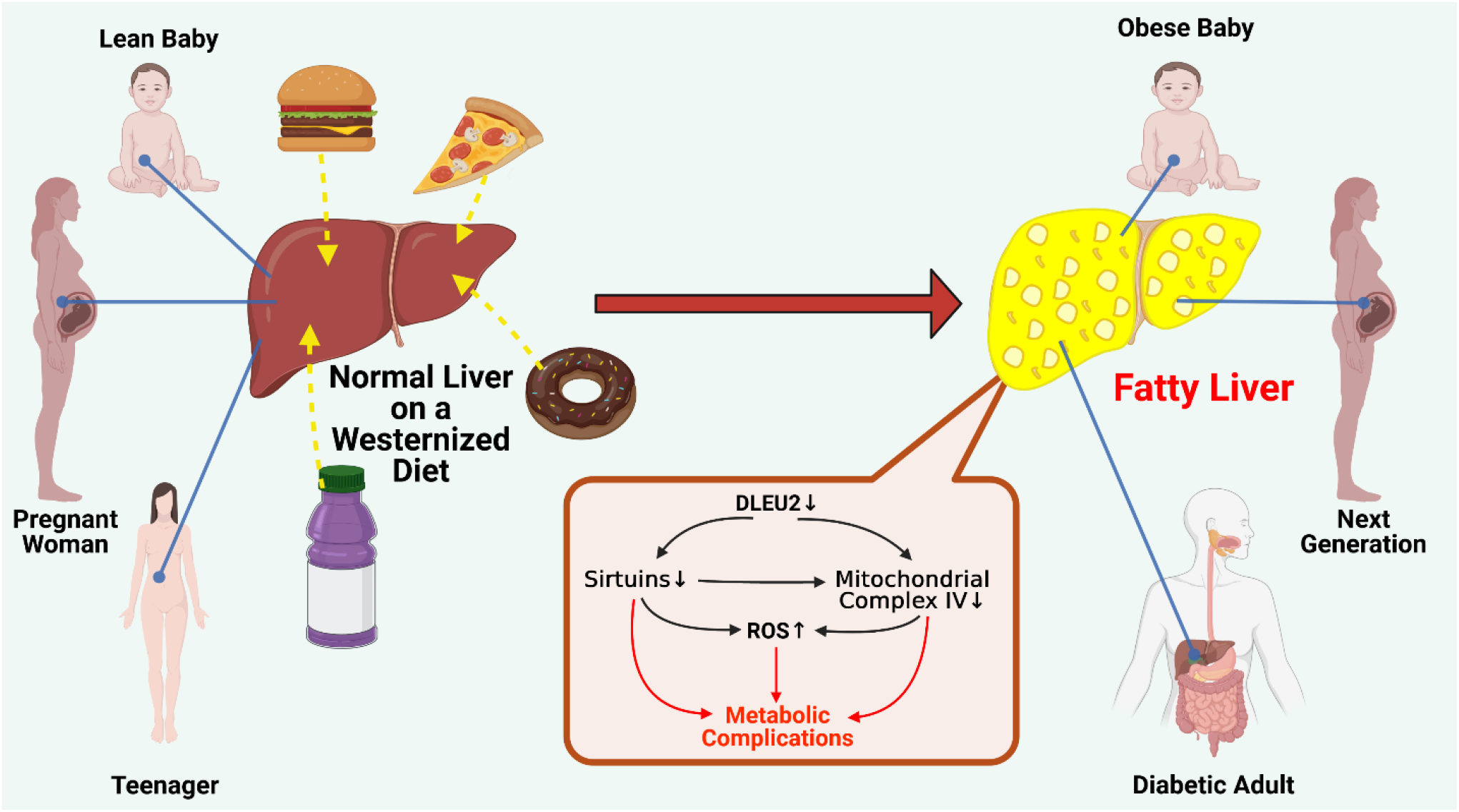
Proposed model of HFD-HF leading to lncRNA DLEU2 regulated epigenetic circuit causing metabolic abnormalities in the liver. Western diet shows effects on DLEU2 induced metabolic regulation leading to mitochondrial dysfunction and oxidative stress in liver. These abnormalities can lead to metabolic diseases including diabetes, obesity, fatty liver and many more. This figure was created with BioRender.com.

## MATERIAL AND METHODS

### Animal studies

Female C57Bl/6 mice (F0) were either fed an ad libitum Control diet (CD (13.2% fat, 24.6% protein, and 62.1% carbohydrates by calories, PicoLab Rodent Diet 5053; Laboratory Supply, Fort Worth, TX) or high-fat/high-fructose diet (HFD-HF; 45% fat, 20% protein, 17% fructose, 17% sucrose by calories, D15041701, Research Diets) for 6 weeks till birth as previously described (Powell and Choudhury, 2019). The male pups (F1) were fed CD or HFD-HF for 20 weeks. Four groups: CD/CD (CD mothers/CD pups, green), CD/HFD-HF (CD mothers/HFD-HF pups, blue), HFD-HF/CD (HFD-HF mothers/CD pups, purple), and HFD-HF/HFD-HF (HFD-HF mothers/HFD-HF pups, red) were assigned. Body weight of F1 was measured at 20 weeks. Liver was excised, weighed, and flash frozen in liquid nitrogen. Comparison of HFD-HF/HFD-HF vs CD/HFD-HF (red vs blue) or HFD-HF/CD vs CD/CD (purple vs green) indicated maternal diet effect (intergenerational effect), and comparison of HFD-HF/HFD-HF vs HFD-HF/CD (red vs purple) or CD/HFD-HF vs CD/CD (blue vs green) indicated offspring diet effect. All procedures were approved by Texas A&M IACUC.

### Cell culture and siRNA knockdown (KD) experiments

HepG2 cells were purchased from ATCC (Manassas, VA). Cells were cultured in EMEM media supplemented with 10% fetal bovine serum, 100 U/mL penicillin, and 100 mg/mL streptomycin (Gibco, NY). The cells were maintained in a humidified incubator at 37°C under an atmosphere of 5% CO_2_. The negative control (Ambion; n410472) or siRNA DLEU2 (Ambion; n274197) was mixed with Lipofectamine RNAiMAX (Invitrogen) according to the manufacturer’s protocol in serum-free OPTI-MEM before incubation. RNA or protein was extracted after 96h transfection.

### mRNA real-time RT-PCR

Liver or cell RNA was extracted according to the Qiagen miRNeasy Mini Kit (Qiagen, MD, USA) protocol. Reverse transcription (RT) was carried out using the High-Capacity cDNA Reverse Transcription Kit (Life Technologies, NY, USA). One microgram of mRNA was reverse transcribed to cDNA according to the manufacturer’s instructions. The expression levels of gene transcripts were determined using quantitative real-time PCR. The qRT-PCR mix contained optimal concentrations of primers, cDNA and SYBR Green PCR Master Mix (Life Technologies, NY, USA). Relative gene expression was normalized to β-actin and represented as fold change (Meruvu et al., 2016b). Primers’ sequences are shown in table 1.

**Table1.**
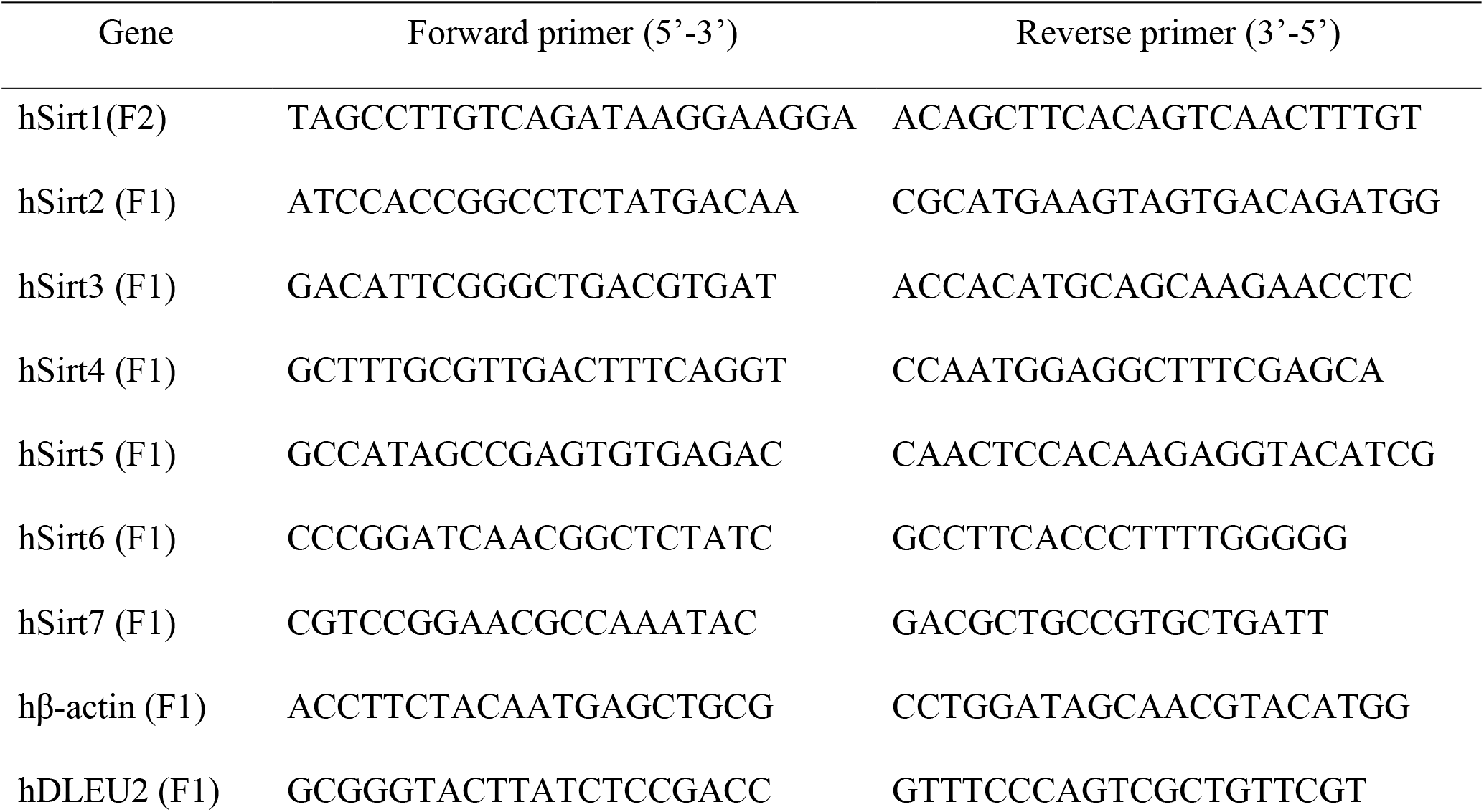
Gene primers used in this study.

### Western blot analysis

Cell were lysed with RIPA lysis buffer (Cat#9803, Cell signaling) plus 0.5 mM PMSF and Protein inhibitor tablets (Cat#A32963, Pierce™, Thermo Scientific). The protein content of the lysate was determined using Bio-Rad Protein Assay (Bio-Rad) on a spectrophotometer. The same amount of isolated proteins were separated by SDS-PAGE in LI-COR running buffer with reduced reagent, and transferred to PVDF membrane. After the membrane was blocked with LI-COR blocking buffer, it was immunoblotted with primary antibody, including SIRT1 (Cat#2028), SIRT2 (Cat#D4S6J), SIRT3 (Cat#5490), SIRT5 (Cat#8779), SIRT6 (Cat#12486), SIRT7 (Cat#5360) or GAPDH (Cat#2118S) from Cell Signaling Technologies, or SIRT4 (ab124521) and Total OXPHOS Rodent WB Antibody Cocktail (ab110413) from Abcam, or β-actin (Cat#PA5-59497) and MTCO1 (Cat#459600) from ThermoFisher Scientific. Finally, Goat anti-rabbit or anti-mouse IRDye 680 or IRDye 800 secondary antibodies were used for the detection and quantitation of immunoblots. Membranes were imaged using a LI-COR Odyssey scanner, and blots were analyzed by Image Studio Lite 5.0 analytical software (LI-COR, Lincoln, NE) as previously described (Meruvu et al., 2016a; Zhang and Choudhury, 2017).

### Oxidative stress assay

Reactive oxygen species (ROS) levels were measured using 2’, 7’-dichlorofluorescein diacetate (DCFDA) as substrate according to the manufacturer’s instructions (Meruvu et al., 2016b; Zhang et al., 2015). Briefly, HepG2 cells were seeded in a 96-well plate knocked down by DLEU2 SiRNA for 96 h as described earlier. Cells were then incubated with 25 μM DCFDA for 45 min at 37 °C in dark conditions, and fluorescence increment was measured for 30 min using a SpectraMax microplate reader (Molecular Devices, CA, USA) at 485 nm excitation and 535 nm emission.

### Statistical analysis

All data are presented as the mean ± standard error (S.E.M). Comparison between groups was performed by using One-way ANOVA (> two groups, followed by Tukey’s test) or Student’s t-test (two groups). A P-value less than 0.05 was considered a statistically significant difference. Statistical analysis was performed using Prism 6.0.

## ACKNOWLEDGEMENT

The work was supported by Morris L Lichtenstein Jr. Medical Research Foundation. We would like to acknowledge Dr. Catherine Powell, for collecting data for body weight and Dr. Raktima Bhattacharya for running some qPCR plates. We also thank Faith Upton for helping the model drawing.

## AUTHOR CONTRIBUTIONS

All three authors intellectually put their effort to hypothesize, carry out, and interpreting the data. M. C. designed the experiments in coordination with J.Z. J. Z. and M.K performed of the experiments, M. K. wrote the manuscript with M. C. and J. Z.

